# Time intervals in sequence sampling, not data modifications, have a major impact on estimates of HIV escape rates

**DOI:** 10.1101/221812

**Authors:** Vitaly V. Ganusov

## Abstract

The ability of HIV to avoid recognition by humoral and cellular immunity (viral escape) is well documented but the strength of the immune response needed to cause such a viral escape remains poorly quantified. Several previous studies observed a more rapid escape of HIV from CD8 T cell responses in the acute phase of infection as compared to the chronic infection. With the help of simple mathematical models the rate of HIV escape was estimated and results were interpreted to suggest that CD8 T cell responses causing escape in acute HIV infection may be more efficient at killing virus-infected cells than responses that cause escape in chronic infection, or alternatively, early escapes occur in epitopes mutations in which there is minimal fitness cost to the virus. These conclusions, however, were challenged on several grounds, including linkage and interference of multiple escape mutations due to a low population size and because of potential issues associated with modifying the data to estimate escape rates. Here we use a parametric resampling method which does not require data modification to show that previous results on the decline of the viral escape rate with time since infection remain unchanged. However, using this method we also show that estimates of the escape rate are highly sensitive to the time interval between measurements with longer intervals biasing estimates of the escape rate downwards. Our results thus suggest that data modifications for early and late escapes were not the primary reason for the observed decline in the escape rate with time since infection. However, longer sampling periods for escapes in chronic infection strongly influence estimates of the escape rate. More frequent sampling of viral sequences in the chronic infection may improve our understanding of factors influencing the rate of HIV escape from CD8 T cell responses.

## Introduction

Due to the absence of a proof-reading capability, HIV-1 (or HIV thereafter) has a relatively high average mutation rate of 2 – 4 × 10^−5^ per bp per replication cycle [1–5]; however, even higher mutation rates have been suggested [5, 6]. Given a relatively short genome (~ 10^4^ bp), mutant viruses arise only in 20–40% of replication cycles. Even with such perhaps surprising fidelity given a short virus replication cycle time (~ 1 – 2 days, [7]) and large population size (~ 10^7^ – 10^8^ of virus-infected cells, [8, 9]), HIV is able to generate multiple variants during the course of infection. Some of these variants will be unrecognized by the host’s immune responses resulting in escape, which may prevent viral control. Indeed, the ability of HIV to escape from cellular (CD8 T cell) and humoral (antibody) responses is often defined as the major obstacle preventing the development of efficient vaccines [10–13]. One of the early mathematical models of why HIV-infected individuals progress to AIDS also involves viral escape from immunity [14, 15].

While HIV clearly escapes from both cellular and humoral immune responses, careful tracking of the escape dynamics suggested that escape from virus-specific CD8 T cell responses occurs earlier than from antibody-mediated responses [16–19]. Escape from CD8 T cell responses has been accurately mapped for several HIV-infected individuals from the earliest stages of infection [16, 19, 20]. Specifically, these studies tracked changes in HIV genome from the virus sequence founding the infection (transmitted/founder virus) and mapped CD8 T cell response specific to whole viral proteome. Genetic changes in regions, recognized by CD8 T cells, was treated as evidence of T cell recognition of the virus and thus, the speed at which the founder viral sequence was lost was indicative of T cell response strength. Multiple studies developed mathematical models to estimate the rate at which founding, wild-type virus was lost from the population (and consequently, the escape variants accumulated in the population); this escape rate was then interpreted as the rate at which epitope-specific CD8 T cells eliminate HIV-infected cells [21–27]. Estimation of HIV escape rates was deemed complicated due to the sparse nature of the sequence data coming from relatively infrequent sampling of virus populations and a low number of viral sequences obtained at a given time point in most studies. In particular, many of the data on escape measuring frequency of the wild-type (or escape) variant include measurements of only either 100% or 0% in two sequential time points (or in some cases, with one intermediate frequency of the wild-type virus in between of the two extreme values of 100% and 0%). To estimate HIV escape rates in such data, the data were modified to include some presence of the escape variant or the wild-type virus [21, 22]. For example, if out of 10 sequences of a specific epitope region 10 sequences were of the wild-type virus, it was proposed to add 1 extra sequence with a mutated epitope resulting in the frequency of the wild-type *f_w_* = 10/11 = 91% instead of the observed 100%. Similarly, if 5 out 5 sequences at the subsequent time point were mutated, adding 1 wild-type sequence resulted in the wild-type variant frequency of *f_w_* = 1/6 = 17% instead of 0%. Such modifications were proposed to provide minimal escape rates [21, 25]. It remains unknown whether such data modifications were important for making general observations regarding HIV escape rates; previous studies utilizing such data modifications suggested that the rate of HIV escape declines with time since infection suggesting potential weakening of the immune responses or a simple consequence of the virus escaping in least costly positions [25, 28]. More recent studies highlighted that the proposed data modification may have had a strong impact on estimated escape rates, thus questioning the validity of previous conclusions [29, 30]. Here we extend previous analyses by applying a novel method of estimating rates of viral escape from poorly sampled data which does not involve modification of the data. The method uses resampling of the sequence data using the binomial distribution. With this resampling method we show that at least for three HIV-infected patients, previous data modifications were not responsible for the predicted decline of the escape rate with time since infection. However, the resampling method remains highly sensitive to the time frequency of data sampling — the method estimates slower escapes for less frequently sampled data. This analysis suggests that better understanding of the mechanisms behind HIV escape from CD8 T cell responses is not likely to come from improved methods of sequence data analysis but from better data with improved time resolution and depth of sequencing.

## Results

### A resampling method to estimate escape rates without data modification

Several previous studies have shown that the kinetics of viral escape from a single CD8 T cell response can be described by the logistic equation

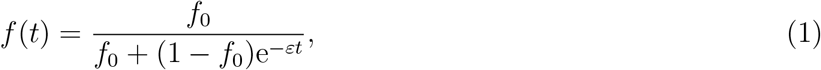

where *f*_0_ and *f*(*t*) is the frequency of the escape variant at time *t* = 0 and time *t*, respectively, and ε is the escape rate [21–23, 25]. When the escape variant is generated by mutation from the wild type, the predicted dynamics of the frequency of the escape variant is approximately given by the logistic equation [31, eqn. (1)]. The time, at which the wild-type or mutant variant reaches 50% is called 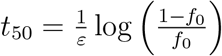[25].

Data on viral escape are generally given as the percent of escape variant in the viral population at different time points with a number of viral sequences obtained. For example, at time 10 days post symptoms, 15 viral sequences were obtained and 3 of these contained mutated epitopes, so the frequency of the escape variant is 3/15 or 20%. To estimate escape rate it is advised to use a maximum likelihood approach [31] in which the binomial distribution-based likelihood of the model given experimental data is given by

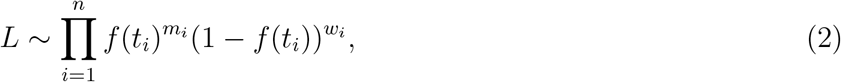

where *m_i_* and *w_i_* are the number of escape mutant and wild-type sequences in the *i^th^* sample taken at time *t_i_* and there are *n* time points. Note that we ignored constant terms for the binomial distribution since these do not matter during likelihood maximization. Parameters of the model including the escape rate ε and initial variant frequency *f*_0_ are then found by maximizing likelihood [31]. As discussed in the introduction, the likelihood function (eqn. (2)) fails if the data do not have measurements in which both wild-type and mutant sequences are observed. In some cases when there is only one time point which both wild-type and mutant viruses are detected and for all other time points only one variant is observed, it is possible to estimate the escape rate without data modification [?].

To obtain an estimate of the escape rate for the data when there is only wild-type or only mutant variants are detected at a given time point we propose to use a parameteric bootstrap. In this approach we do not fit the basic mathematical model (eqn. (1)) to the data. Instead, we generate pseudorandom samples of the data which are found using a beta distribution for every sample in every time point [32]. Specifically, for the *N_i_* = *w_i_* + *m_i_* sampled sequences at time *t_i_* where *m_i_* is the number of escape variant sequences and *w_i_* is the number of wild-type sequences, the resampled frequency of the escape variant *x_i_* is given by the beta distribution *Beta*(*m_i_* + 1/2, *w_i_* + 1/2) [32]. This procedure is repeated for all *n* time points resulting in a bootstrap sample. In our analyses we used the routine `Random` in `Mathematica` (the Mathematica code providing the method is available as a supplement to this article). Due to randomness, the resulting bootstrapped sample will have both wild-type and escape variants present at multiple time points. The model then can be fitted to these bootstrapped data using the likelihood approach (eqn. (2)). Repeating this sampling procedure a given number of times (e.g., 10^3^ simulations), a distribution of escape rates, initial mutant frequency, and the time to 50% of the escape variant can be generated. The mean (or median) of these distributions and 95% interval of the distribution can be then used to characterize the escape rate ε and the time to 50% escape variant *t*_50_.

While the resampling method was specifically designed to estimate escape rates from the data which cannot be fitted by simple mathematical models (e.g., as in eqn. (1)), it was important to confirm that for well sampled data the method is able to recover the escape rate that would be found by fitting the models directly to such data. As a proof of principle we simulated viral escape using the logistic model (eqn. (1)) and assumed that it was possible to detect both wild-type and escape variant viruses at three sequential time points (Figure 1). Obviously, the logistic model can well fit these data (Figure 1A) without the need for additional methods. Importantly, however, resampling these data using the beta distribution and estimating the escape rate for these bootstrap samples allowed for a relatively accurate estimate of the fast and slow escape rates (Figure 1C). Additional simulations with more randomly chosen escape rates and samplings confirmed that the method of resampling recovers the same escape as the model fitted directly to the escape data. The results of these analyses were similar if we used data in which both wild-type and escape variants were present at 2 sequential time points (results not shown). In our simulations, resampling using beta distribution never yielded a 0% or 100% for the wild-type virus, and thus always allowed us to find a finite value for the escape rate.

**Figure 1:**
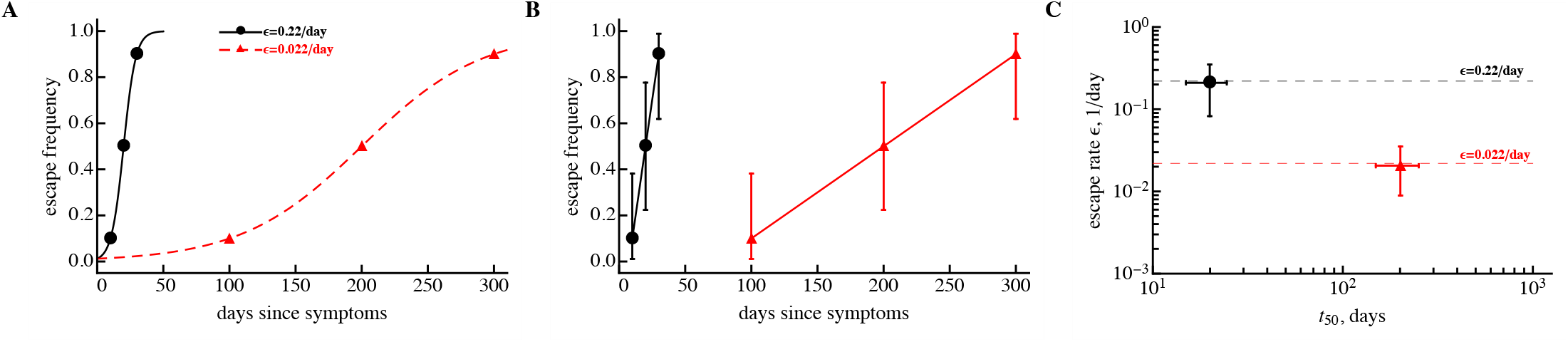
The resampling (parametric bootstrap) method accurately recovers virus escape rate for data when both wild-type and escape variant are detected at multiple sequential time points. We simulated viral escape using a logistic equation with two escape rates: *ε* = 0.22/day (bullets) and *ε* = 0.022/day (triangles). Three data points at which the escape variant was present at 10%, 50%, and 90% out of N = 10 sequences were generated (panel A). The data were fitted by the simple logistic model (eqn. (1)) using a likelihood approach leading to expected escape rates (lines in panel A). These data were then resampled using the beta distribution (panel B) and fitted by the same model using the likelihood method. Resulting averages and their 95% confidence intervals match well to the numbers estimated using actual data (panel C). Dashed lines in panel C show estimates of the escape rates found in panel A. 95% confidence intervals in panel B are predicted using Jeffrey’s intervals [32].

### The rate of HIV escape declines with time since infection

Previous work from several groups suggested that during HIV infection the rate of HIV escapes from T cell responses declines over time [25, 28]. However, this conclusion was challenged on multiple levels including the need for modification of the data to provide minimal estimates of the escape rate [29]. Indeed, it was possible that for many late escapes previous studies provided minimal estimates of the escape rate and thus the observed pattern of the escape rate decline with time since infection was an artifact of data modification [26, 29, 30]. To address this specific point we re-analyzed data on HIV escape published previously [16, 25]. Specifically, we used data for all documented escapes in three patients from the Center for HIV/AIDS Vaccine Immunology (CHAVI) as described in previous publications and limited these data to the first 800 days post-infection. This was done because of the bias towards lower estimates the resampling method introduces in datasets with longer sampling times (see Discussion). We then resampled data for all escapes, fitted the simple logistic model (eqn. (1)) to these data and calculated the average escape rate (ε) and time of escape (*t*_50_). This reanalysis revealed a consistent, statistically significant decline in the estimated rate of HIV escape from CD8 T cell responses with the time since infection (Figure 2). This was true for all documented escapes (i.e., escapes which had the pattern of viral escape from T cell responses but may or may not have had measurable T cell response against the wild-type epitope) in 2 out of 3 patients with escapes with detected T cell responses against the wild-type epitope. The poor correlation in the latter case in patient CH58 is likely due to a small number of confirmed escapes (*n* = 4). Interestingly, there was a strong correlation between mean escape rates obtained from parametric resampling of the data and previous estimates found with data modifications (results not shown). Thus, this analysis suggests that data modification was not the reason for the detected decline in the escape rate with time since infection [25].

**Figure 2:**
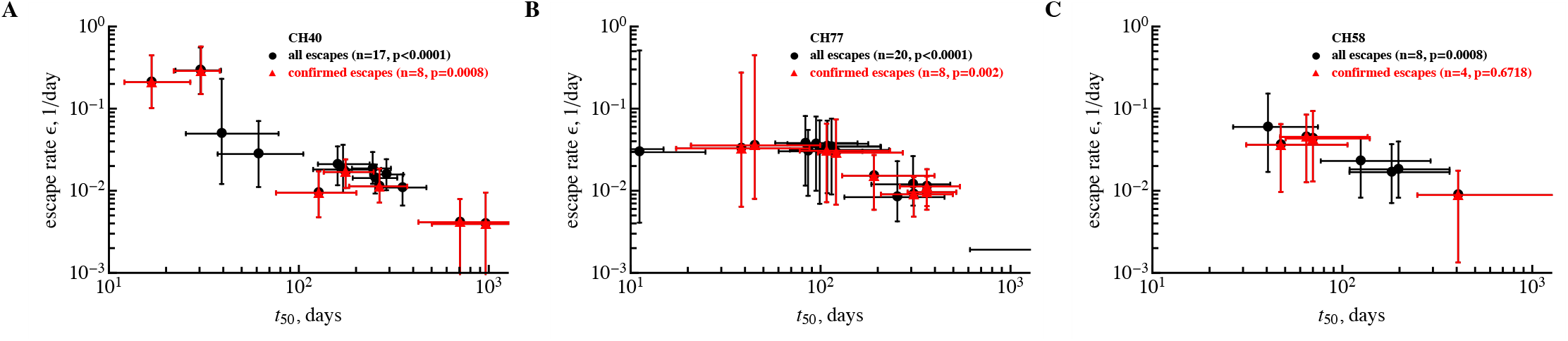
A resampling method which does not involve data modification still predicts a decline in the estimated escape rate with the time since infection. We used a novel approach of data resampling and estimated the rate of HIV escape from CD8 T cell responses for three CHAVI patients using previously published data [16, 25]. Rates (*ε*) and time to 50% of the escape variant (*t*_50_) and associated errors are shown for 3 different patients in different panels (CH40, CH77, CH58); bullets indicate all escapes (in black) and triangles are for confirmed escapes (in red) [16]. Confirmed escapes were defined as fixed changes in HIV genome with a clear signature of viral escape with detected epitope-specific T cell response. Other escapes had a well defined signature of escape but no epitope-specific T cell response was detected [16]. Resulting p values from the Spearman Rank correlation test are indicated on individual panels. Panel B does not show an estimate for one escape due to very low escape rate *ε* and long *t*_50_.

### Time frequency of sampling biases estimates of the escape rate

Our results so far suggest that data modifications were not likely responsible for the observed decline in the estimated escape rate with time since infection. Therefore, it is important to understand whether other features in the data could be responsible for this declining trend. Previous studies have highlighted that the frequency of sampling in experimental data have changed over time starting from weekly sampling and changing to monthly/bi-monthly sampling [16]. Such a pattern may naturally lead to underestimates of escape rates [26, 29, 30]. Therefore, we sought to determine if the resampling method is capable of accurately recovering the escape rate independently of the sampling time period. We simulated viral escape with the same escape rate but assumed different initial frequency for the virus and different time intervals between measurements. This allowed wild-type and escape variant to be present in the data only at one time point (Figure 3A). The resulting “data” looked somewhat similar to those observed during HIV infection (Figure 3B). By fitting the logistic model to bootstrapped resamples we found that the sampling period has a dramatic impact on the estimated escape rate (Figure 3C). In an extreme case the method underestimates the escape rate 10 fold! Thus, time frequency of sampling (i.e., weekly, monthly, bi-monthy, etc.) has a dramatic impact on the estimate of the escape rate even in the absence of data modification and when one uses a resampling method. This result has two consequences. First, in order to understand the kinetics of HIV escape from T cell responses more frequent sampling during chronic HIV infection will be needed. Second, previous results on the independence of the escape rate on the magnitude of the T cell response [25] and the decline of the escape rate with the time since infection have to be re-evaluated using additional, perhaps better sampled data.

**Figure 3:**
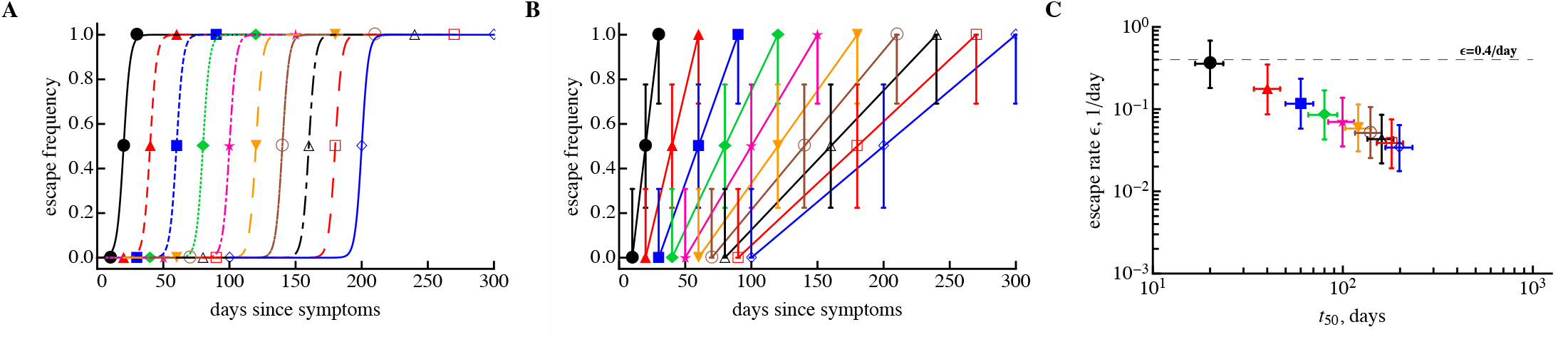
Time interval between measurements has a strong bias on the estimate of the escape rate and leads to a correlation between the escape rate (*ε*) and time of escape (*t*_50_). We generated artificial data in which viral sequences were measured in three time points, resulting in 0%, 50%, and 100% of the escape variant with *N* = 10 sequences per time point (panel A). In all cases, escape can be described by the logistic equation (eqn. (1)) assuming delayed immune response and escape rate *ε* = 0.4/day. We resampled these simulated “data” using a beta distribution (panel B) and estimated the average escape rate (panel C). We show the prediction of the logistic curve obtained using *ε* = 0.4/day (panel A), sampled data (panel B) and estimates of the escape rate obtained using parametric bootstrap (panel C). The horizontal dashed line in panel C denotes the theoretical escape rate of 0.4/day. The 95% confidence intervals in panel B are predicted using Jeffrey’s intervals [32] and in panel C were calculated from bootstrapped samples.

## Discussion

CD8 T cell responses have been hypothesized to play an important role in control of HIV replication; however, for humans, direct evidence of such a role is lacking. Escape of HIV from T cell responses has been cited as one type of indirect evidence in support of this hypothesis. Accurate estimates of the rate at which CD8 T cells are eliminating the virus-infected cells (or more generally, in suppressing virus replication) and comparison of these rates with other kinetic parameters of HIV replication *in vivo* may be useful in testing this hypothesis. The rate at which HIV escapes from CD8 T cell responses may be one metric indicating the strength of the HIV-specific CD8 T cell response. However, multiple studies have now shown that estimating the rate of HIV escape from current data is complicated.

We have proposed a new method of resampling sequence data to estimate escape rates in situations when direct fitting of simple mathematical models to the data fails. An alternative approach in estimating escape rates from sparsely sampled data would be to use more mechanistic models of HIV dynamics and evolution, for example, models that take into account virus effective population size, the rate of HIV mutation and recombination [26]. However, it is unclear how well such complex models with multiple parameters represent HIV dynamics *in vivo*, and as such, biases in parameter estimates associated with such models are not known.

Using the resampling method we have shown that the observed phenomenon of a decline of the rate of HIV escape with time since infection is not the result of modification of data to accommodate fits of simple models of viral escape. Yet, the method suffers from a great sensitivity to the sampling frequency in the data since increasing the time intervals between measurements naturally decreases the estimated escape rate. Furthermore, inclusion of more data at later time points in which wild-type virus is nearly lost will also bias average estimates found by parameteric bootstrap downwards which further limits the applicability of this method for estimation of escape rates (results not shown). Specifically, if for the first escape in Figure 1A we added more measurements at later time points, say at day 100 or 300 (when the escape variant is likely to be at 100%), resampling would have provided much lower estimates of the escape rate than the ε = 0.22/day found by the model fitting to original, not resampled data (results not shown). The resampling method also requires more computational power as at least 10^3^ samples are generally required to accurately estimate confidence intervals [33]. While fitting a simple logistic model to escape data is generally fast, fitting more complicated models assuming, for example, sequential or concurrent escape, can be time consuming [34]. Finally, the method relies on the approximation of the binomial distribution by the beta distribution which may fail in some circumstances [32].

Overall, our results so far suggest that the current state of data on HIV escape does not allow us to make solid inferences on the escape rate without making strong assumptions either about the data or about the underlying dynamics of the virus. It appears that the way forward is to generate novel data in which patient sampling occurs at regular intervals over the course of a few months of infection perhaps with the use of novel techniques such as deep sequencing. Consistent temporal sampling rather than deep sampling is likely to be more beneficial at determining if the rate of HIV escape from T cell responses declines with time since infection [31].

## References

1. Mansky, L. M. and H. M. Temin. 1995. Lower in vivo mutation rate of human immunodeficiency virus type 1 than that predicted from the fidelity of purified reverse transcriptase. J Virol 69: 5087–94.

2. Mansky, L. M. 1996. Forward mutation rate of human immunodeficiency virus type 1 in a T lymphoid cell line. AIDS Res Hum Retroviruses 12: 307–14.

3. Mansky, L. M. 1998. Retrovirus mutation rates and their role in genetic variation. J Gen Virol 79: 1337–45.

4. Sanjun, R., M. R. Nebot, N. Chirico, L. M. Mansky, and R. Belshaw. 2010. Viral mutation rates. J Virol 84: 9733–9748.

5. Geller, R., P. Domingo-Calap, J. M. Cuevas, P. Rossolillo, M. Negroni, and R. Sanjun. 2015. The external domains of the hiv-1 envelope are a mutational cold spot. Nature communications 6: 8571.

6. Cuevas, J. M., R. Geller, R. Garijo, J. Lpez-Aldeguer, and R. Sanjun. 2015. Extremely high mutation rate of hiv-1 in vivo. PLoS biology 13: e1002251.

7. Perelson, A. S. 2002. Modelling viral and immune system dynamics. Nature Rev Immunol 2: 28–36.

8. Haase, A. T. 1999. Population biology of HIV-1 infection: viral and CD4+ T cell demographics and dynamics in lymphatic tissues. Annu Rev Immunol 17: 625–56.

9. Estes, J. D., C. Kityo, F. Ssali, L. Swainson, K. N. Makamdop, G. Q. Del Prete, S. G. Deeks, P. A. Luciw, J. G. Chipman, G. J. Beilman, T. Hoskuldsson, A. Khoruts, J. Anderson, C. Deleage, J. Jasurda, T. E. Schmidt, M. Hafertepe, S. P. Callisto, H. Pearson, T. Reimann, J. Schuster, J. Schoephoerster, P. Southern, K. Perkey, L. Shang, S. W. Wietgrefe, C. V. Fletcher, J. D. Lifson, D. C. Douek, J. M. McCune, A. T. Haase, and T. W. Schacker. 2017. Defining total-body aids-virus burden with implications for curative strategies. Nature medicine 23: 1271–1276.

10. Walker, B. D. and B. T. Korber. 2001. Immune control of HIV: the obstacles of HLA and viral diversity. Nature immunology 2: 473–475.

11. Barouch, D., J. Kunstman, M. Kuroda, J. Schmitz, S. Santra, F. Peyerl, G. Krivulka, K. Beaudry, M. Lifton, D. Gorgone, D. Montefiori, M. Lewis, S. Wolinsky, and N. Letvin. 2002. Eventual AIDS vaccine failure in a rhesus monkey by viral escape from cytotoxic T lymphocytes. Nature 415: 335–9.

12. Mullins, J. I., M. Rolland, and T. M. Allen. 2008. Viral evolution and escape during primary human immunodeficiency virus-1 infection: implications for vaccine design. Curr Opin HIV AIDS 3: 60–66.

13. Rolland, M., S. Tovanabutra, A. C. Decamp, N. Frahm, P. B. Gilbert, E. Sanders-Buell, L. Heath, C. A. Magaret, M. Bose, A. Bradfield, A. O’Sullivan, J. Crossler, T. Jones, M. Nau, K. Wong, H. Zhao, D. N. Raugi, S. Sorensen, J. N. Stoddard, B. S. Maust, W. Deng, J. Hural, S. Dubey, N. L. Michael, J. Shiver, L. Corey, F. Li, S. G. Self, J. Kim, S. Buchbinder, D. R. Casimiro, M. N. Robertson, A. Duerr, M. J. McElrath, F. E. McCutchan, and J. I. Mullins. 2011. Genetic impact of vaccination on breakthrough HIV-1 sequences from the STEP trial. Nat Med 17: 366–71.

14. Nowak, M. A., R. M. Anderson, A. R. McLean, T. F. Wolfs, J. Goudsmit, and R. M. May. 1991. Antigenic diversity thresholds and the development of AIDS. Science 254: 963–9.

15. Alizon, S. and C. Magnus. 2012. Modelling the course of an hiv infection: insights from ecology and evolution. Viruses 4: 1984–2013.

16. Goonetilleke, N., M. K. Liu, J. F. Salazar-Gonzalez, G. Ferrari, E. Giorgi, V. V. Ganusov, B. F. Keele, G. H. Learn, E. L. Turnbull, M. G. Salazar, K. J. Weinhold, S. Moore, N. Letvin, B. F. Haynes, M. S. Cohen, P. Hraber, T. Bhattacharya, P. Borrow, A. S. Perelson, B. H. Hahn, G. M. Shaw, B. T. Korber, and A. J. McMichael. 2009. The first T cell response to transmitted/founder virus contributes to the control of acute viremia in HIV-1 infection. J Exp Med 206: 1253–72.

17. McMichael, A. J., P. Borrow, G. D. Tomaras, N. Goonetilleke, and B. F. Haynes. 2010. The immune response during acute HIV-1 infection: clues for vaccine development. Nat Rev Immunol 10: 11–23.

18. Bar, K. J., C.-Y. Tsao, S. S. Iyer, J. M. Decker, Y. Yang, M. Bonsignori, X. Chen, K.-K. Hwang, D. C. Montefiori, H.-X. Liao, P. Hraber, W. Fischer, H. Li, S. Wang, S. Sterrett, B. F. Keele, V. V. Ganusov, A. S. Perelson, B. T. Korber, I. Georgiev, J. S. McLellan, J. W. Pavlicek, F. Gao, B. F. Haynes, B. H. Hahn, P. D. Kwong, and G. M. Shaw. 2012. Early Low-Titer Neutralizing Antibodies Impede HIV-1 Replication and Select for Virus Escape. PLoS Pathog 8: e1002721.

19. Kijak, G. H., E. Sanders-Buell, A.-L. Chenine, M. A. Eller, N. Goonetilleke, R. Thomas, S. Leviyang, E. A. Harbolick, M. Bose, P. Pham, C. Oropeza, K. Poltavee, A. M. O’Sullivan, E. Billings, M. Merbah, M. C. Costanzo, J. A. Warren, B. Slike, H. Li, K. K. Peachman, W. Fischer, F. Gao, C. Cicala, J. Arthos, L. A. Eller, R. J. O’Connell, S. Sinei, L. Maganga, H. Kibuuka, S. Nitayaphan, M. Rao, M. A. Marovich, S. J. Krebs, M. Rolland, B. T. Korber, G. M. Shaw, N. L. Michael, M. L. Robb, S. Tovanabutra, and J. H. Kim. 2017. Rare hiv-1 transmitted/founder lineages identified by deep viral sequencing contribute to rapid shifts in dominant quasispecies during acute and early infection. PLoS pathogens 13: e1006510.

20. Liu, M. K. P., N. Hawkins, A. J. Ritchie, V. V. Ganusov, V. Whale, S. Brackenridge, H. Li, J. W. Pavlicek, F. Cai, M. Rose-Abrahams, F. Treurnicht, P. Hraber, C. Riou, C. Gray, G. Ferrari, R. Tanner, L.-H. Ping, J. A. Anderson, R. Swanstrom, M. Cohen, S. S. A. Karim, B. Haynes, P. Borrow, A. S. Perelson, G. M. Shaw, B. H. Hahn, C. Williamson, B. T. Korber, F. Gao, S. Self, A. McMichael, and N. Goonetilleke. 2013. Vertical T cell immunodominance and epitope entropy determine HIV-1 escape. J Clin Invest 123: 380–393.

21. Fernandez, C., I. Stratov, R. De Rose, K. Walsh, C. Dale, M. Smith, M. Agy, S. Hu, K. Krebs, D. Watkins, D. O’connor, M. Davenport, and S. Kent. 2005. Rapid viral escape at an immunodominant simian-human immunodeficiency virus cytotoxic T-lymphocyte epitope exacts a dramatic fitness cost. J Virol 79: 5721–31.

22. Asquith, B., C. Edwards, M. Lipsitch, and A. McLean. 2006. Inefficient cytotoxic T lymphocyte-mediated killing of HIV-1-infected cells in vivo. PLoS Biol 4: e90.

23. Ganusov, V. V. and R. J. De Boer. 2006. Estimating costs and benefits of CTL escape mutations in SIV/HIV Infection. PLoS Comput Biol 2: e24.

24. Fischer, W., V. V. Ganusov, E. E. Giorgi, P. T. Hraber, B. F. Keele, T. Leitner, C. S. Han, C. D. Gleasner, L. Green, C. C. Lo, A. Nag, T. C. Wallstrom, S. Wang, A. J. McMichael, B. F. Haynes, B. H. Hahn, A. S. Perelson, P. Borrow, G. M. Shaw, T. Bhattacharya, and B. T. Korber. 2010. Transmission of single HIV-1 genomes and dynamics of early immune escape revealed by ultra-deep sequencing. PLoS One 5: e12303.

25. Ganusov, V. V., N. Goonetilleke, M. K. P. Liu, G. Ferrari, G. M. Shaw, A. J. McMichael, P. Borrow, B. T. Korber, and A. S. Perelson. 2011. Fitness Costs and Diversity of the Cytotoxic T Lymphocyte (CTL) Response Determine the Rate of CTL Escape during Acute and Chronic Phases of HIV Infection. J Virol 85: 10518–10528.

26. Kessinger, T. A., A. S. Perelson, and R. A. Neher. 2013. Inferring HIV Escape Rates from Multi-Locus Genotype Data. Front Immunol 4: 252.

27. Leviyang, S. 2014. Constructing lower-bounds for ctl escape rates in early siv infection. Journal of theoretical biology 352: 82–91.

28. Asquith, B. and A. McLean. 2007. In vivo CD8+ T cell control of immunodeficiency virus infection in humans and macaques. Proc Natl Acad Sci USA 104: 6365–70.

29. Leviyang, S. and V. V. Ganusov. 2015. Broad CTL Response in Early HIV Infection Drives Multiple Concurrent CTL Escapes. PLoS Comput Biol 11: e1004492.

30. Garcia, V., M. W. Feldman, and R. R. Regoes. 2016. Investigating the Consequences of Interference between Multiple CD8+ T Cell Escape Mutations in Early HIV Infection. PLoS Comput Biol 12: e1004721.

31. Ganusov, V. V., R. A. Neher, and A. S. Perelson. 2013. Modeling HIV escape from cytotoxic T lymphocyte responses. J Stat Mech 2013: P01010.

32. Brown, L., T. Cai, A. DasGupta, A. Agresti, B. Coull, G. Casella, C. Corcoran, C. Mehta, M. Ghosh, T. Santner, L. Brown, T. Cai, and A. DasGupta. 2001. Interval estimation for a binomial proportion. Statistical Science 16: 101–133.

33. Efron, B. and R. Tibshirani. 1993. An introduction to the bootstrap. Chapman & Hall, New York, 436 p.

34. Yang, Y. and V. V. Ganusov. 2017. Kinetics of HIV-specific CTL responses plays a minimal role in determining HIV escape dynamics. Front Immunol (in review).

